# Toward routine health phenotyping: High-throughput prediction of metabolic, immune, and inflammatory biomarkers from milk mid-infrared spectroscopy in early-lactation dairy cows

**DOI:** 10.64898/2026.07.14.738588

**Authors:** P.N. Ho, J.E. Hemsworth, C.M. Reich, C.R. Bath, Z. Liu, S. Rochfort, M. Khansefid, S.T. Muhammad, M. Haile-Mariam, M.E. Goddard, L.C. Marett, S.R.O. Williams, C.K.M. Ho, M. J. Berkhout, R. Xiang, A.J. Chamberlain

**Affiliations:** Agriculture Victoria, AgriBio, Centre for AgriBioscience, Bundoora, Victoria 3083, Australia; Agriculture Victoria, Ellinbank, Victoria 3821, Australia; School of Applied Systems Biology, La Trobe University, Bundoora, Victoria 3083, Australia; School of Agriculture, Food and Ecosystem Sciences, Faculty of Science, the University of Melbourne, Victoria 3010 Australia

**Keywords:** dairy cows, metabolic disorders, serum biomarkers, mid-infrared spectroscopy

## Abstract

This study evaluated the potential of milk mid-infrared (MIR) spectroscopy, combined with routinely available on-farm variables, for predicting serum metabolic, immune, and inflammatory biomarkers in early-lactation cows. Data included 5,936 blood samples from 4,442 cows across 23 Australian dairy herds, with paired milk MIR spectra and serum measurements for up to 14 biomarkers. Prediction models were developed using partial least squares regression and evaluated using nested 10-fold random cross-validation and leave-one-herd-out validation. The results show that while basic herd-test data, including milk fat, protein, and lactose concentration, as well as on-farm variables, including DIM, calving age, breed, and herd could predict serum biomarkers, combining MIR spectra with these on-farm variables produced the best overall performance. In random cross-validation, blood urea nitrogen (BUN) was predicted most accurately (R^2^ = 0.78), while β-hydroxybutyrate (BHB) and nonesterified fatty acids (NEFA) showed moderate accuracy (R^2^ = 0.56 and 0.44, respectively). BUN also showed the strongest external validation performance, with leave-one-herd-out R^2^ = 0.58 and comparable accuracy for predicting records collected after 70 days in milk (R^2^ = 0.65). BHB and NEFA had moderate leave-one-herd-out accuracy but did not transfer beyond early lactation. Most other biomarkers showed low or inconsistent external validation performance. Overall, MIR spectroscopy combined with on-farm variables shows promise for routine prediction of BUN, BHB and NEFA, which can be used for monitoring and genetic evaluation of, for example, ketosis and energy deficit. Initial random cross-validation results for glucose, bilirubin and cholesterol were promising, but more data is needed to improve the prediction accuracy and robustness of the predictions.

**Highlights:**

- Milk MIR can predict several serum biomarkers in early-lactation dairy cows.
- MIR-predicted blood urea nitrogen shows the greatest accuracy and robustness.
- MIR-predicted β-hydroxybutyrate and nonesterified fatty acids show moderate accuracy.
- Most mineral, hepatic, and inflammatory biomarkers had limited accuracy.
- Routine MIR phenotyping is most promising for BUN, BHB, and NEFA.

**Summary:** Milk mid-infrared (MIR) spectroscopy and on-farm variables are evaluated as a high-throughput tool to predict health-related serum biomarkers in early-lactation dairy cows. The data include 5,936 paired blood and milk samples from 4,442 cows across 23 Australian dairy herds and up to 14 serum biomarkers. Prediction models are developed using partial least squares regression with nested random cross-validation and leave-one-herd-out validation. Blood urea nitrogen (BUN) shows the greatest and most transferable prediction accuracy across herds and lactation stages. β-hydroxybutyrate (BHB) and nonesterified fatty acids (NEFA) are predicted with moderate accuracy, but only during early lactation. Random cross-validation results for glucose and bilirubin are promising, but larger datasets are needed for robust external validation. Most other mineral, hepatic, and inflammatory biomarkers show limited external prediction accuracy. These results indicate that MIR-based routine health phenotyping is most promising for BUN, BHB, and NEFA.

---

Early postpartum is the most metabolically demanding stage of the dairy cow’s productive life which is characterized by rapid shifts in energy balance, liver function, mineral homeostasis, and immune status following the onset of lactation (Overton et al., 2017, Trevisi et al., 2025). These immense physiological changes increase the risk of metabolic disorders such as ketosis, hypocalcaemia, hypomagnesemia, and fatty liver, with well-documented negative effects on milk yield, health, and reproduction (Kang et al., 2025). The economic impact of impaired transition health is substantial, including increased treatment costs, greater culling risk, and reduced milk production and fertility. Therefore, monitoring metabolic status during early lactation is critical for supporting timely management interventions. In addition, routinely generated metabolic health indicators could provide valuable phenotypes for genomic selection to improve transition cow health (Pryce et al., 2016, Luke et al., 2019a).

Serum metabolic profiling is widely used to assess the metabolic health and nutritional status of dairy cows. Although panels vary, they commonly include β-hydroxybutyrate (BHB) and nonesterified fatty acids (NEFA) as indicators of energy balance and ketosis risk; albumin and blood urea nitrogen (BUN) as indicators of protein metabolism, inflammatory status, and nutritional adequacy; calcium (Ca) and magnesium (Mg) as indicators of macromineral status; and bilirubin, cholesterol, triglycerides, and glucose as measures of hepatic function and lipid and carbohydrate metabolism (Overton et al., 2017). Haptoglobin is also an important acute-phase protein because elevated concentrations reflect systemic inflammation (Lassallette et al., 2026). Routine blood sampling in dairy cows is, however, labor-intensive, costly, and invasive, limiting its feasibility for repeated monitoring at herd scale.

Milk mid-infrared (MIR) spectroscopy has emerged as a promising approach to predict metabolic and health-related biomarkers in dairy cows. Among these, BHB, NEFA, and BUN are the most extensively studied, with reported prediction accuracies (R^2^) from random cross-validation ranging from 0.50 to 0.70, 0.39 to 0.53, and 0.54 to 0.87, respectively (Grelet et al., 2018, Benedet et al., 2019, Ho et al., 2021). Giannuzzi et al. (2023) demonstrated that MIR spectroscopy can predict a broad panel of 29 blood biomarkers (R^2^ = 0.20 - 0.83), including those related to energy metabolism, liver function, oxidative stress, inflammation, and mineral status. Milk MIR offers a practical alternative to direct blood assays because it is rapid, cost-effective, and already implemented routinely by commercial milk-testing organizations for milk component testing (i.e. fat, protein and lactose concentrations). As such, MIR-based prediction models for health-relevant biomarkers could enable population-level phenotyping without additional sampling costs, facilitating both herd management and large-scale genetic evaluation (Luke et al., 2019a, Dale et al., 2021).

Building on our previous Australian work demonstrating the promise of MIR spectroscopy for predicting BHB, NEFA, and BUN (Luke et al., 2019b, Ho et al., 2021), the objective of this study was to extend the analysis to a broader panel of metabolic, immune, and inflammatory biomarkers in early-lactation cows, including Ca, Mg, total protein, albumin, globulin, bilirubin, cholesterol, haptoglobin, triglycerides, and glucose.

The data used in this study consisted of 5,936 blood samples taken from 4,442 lactating cows of 23 dairy herds located in Victoria, New South Wales, and Tasmania (Australia) between July 2017 and June 2024. This dataset also included previous records from Luke et al. (2019b) and Ho et al. (2021). Cows were in their first to ninth parity with 54.5% being Holstein-Friesian, 4.7% purebred Jersey and 40.8% crossbred. DIM varied from 1 to 305, with 89.5% of samples taken within 70 days after calving - the period before the peak milk yield and where the majority of metabolic health problems occur.

The blood samples were analyzed for concentrations of BHB, NEFA, BUN, Ca, Mg, total protein, albumin, globulin, bilirubin, cholesterol, haptoglobin, triglycerides, and glucose. Notably, not all samples were analyzed for all biomarkers. The following assays were used: enzymatic kinetic assays for BHB and BUN; enzymatic end-point assay for NEFA; colorimetric complexometric assays using Arsenazo III (pH 6.5) for calcium and Xylidyl Blue for magnesium; biuret method for total protein; bromocresol green (pH 4.2) method for albumin; diazo colorimetric Jendrassik-Grof method for bilirubin; end-point enzymatic colorimetric assays for cholesterol, triglycerides and glucose; and hemoglobin-binding peroxidase activity colorimetric kinetic assay for haptoglobin. Globulin concentration was calculated as total protein minus albumin and the albumin-to-globulin ratio was also calculated. Serum analyses for samples collected between 2017 and 2020 were conducted at Regional Laboratory Services (Benalla, Victoria, Australia) using a Konelab 20XTi automated clinical chemistry analyzer (Thermo Fisher Scientific, Vantaa, Finland). All samples collected thereafter were analyzed at the AgriBio Laboratory, Agriculture Victoria (Bundoora, Victoria, Australia), using a ChemWell 2910 Automated EIA and Chemistry Analyzer (Awareness Technology, Inc., Palm City, FL, USA), using Catachem Inc. (Oxford, CT, USA) reagents, controls, and calibrations as per the manufacturer’s instructions. The AgriBio instrument was calibrated and validated to be consistent with that of Regional Lab Services.

Milk samples were collected within 12 hours of blood sampling and were analyzed for concentration of fat, protein and lactose, and somatic cell count (SCC), using a NexGen Series FTS Combi instrument (Bentley Instruments, Chaska, MN, USA). All milk samples collected in 2017, together with the ones from two Tasmanian herds in 2018 were processed at TasHerd Pty Ltd. (Hadspen, Tasmania, Australia), whereas the remaining milk samples were processed at Hico Pty Ltd. (Maffra, Victoria, Australia). In addition to milk composition, MIR spectra generated by the instruments were retained and used in this study. Each recorded spectrum comprised 899 data points, with each point representing infrared absorbance at a specific wavenumber within the 649 to 3,999 cm^-1^ region.

All animal procedures in this study were conducted in accordance with the Australian Code of Practice for the Care and Use of Animals for Scientific Purposes (NHMRC, 2013). Ethical approvals were granted by the Department of Energy, Environment and Climate Action (DEECA) Agricultural Research and Extension Animal Ethics Committee, who were licensed to conduct animal research in Victoria, New South Wales and Tasmania for all samples collected after April 2020 and the Tasmanian Department of Primary Industries, Parks, Water and Environment (Animal Biosecurity and Welfare Branch, New Town, Tasmania, Australia) for the two Tasmanian herds collected in 2018.

Prior to model development, several mathematical treatments were applied to the serum biomarkers and spectra. A logarithmic (10) and a square root transformation were applied to the original records of BHB and NEFA, respectively. This was done because their natural distributions were not normal, which might impair the accuracy of predicting high values in partial least squares regression. For the spectra, noisy regions (1,615 - 1,652 cm^-1^ and 649 - 925 cm^-1^) characterized by a low signal-to-noise ratio, which is the consequence of a high-water absorption and noninformative region (2,998 - 3,998 cm^-1^), were first removed. Second, to discard the spectra that are potentially outliers, a standardized Mahalanobis distance between each spectrum and the population average was calculated. Then, the spectra with a global distance greater than 3 (n = 26) were considered outliers and eliminated. Last, a first-order Savitzky-Golay derivative was applied to the reduced spectra. The final spectra used for model development consisted of 536 wavenumbers. We also attempted to standardize spectra generated by the TasHerd instrument to those generated by the HICO instrument using 100 identical milk samples and Piecewise Direct Standardization method (Wang et al., 1991), but this produced results comparable to those obtained using the original spectra. This agrees with Pierre (2017), who showed that the Bentley IR cell and spectra standardization gave consistent BHB predictions over time and globally. Therefore, original spectra were used for all analyses. The number of records and summary statistics for each biomarker are summarized in Table 1.

**Table 1.**
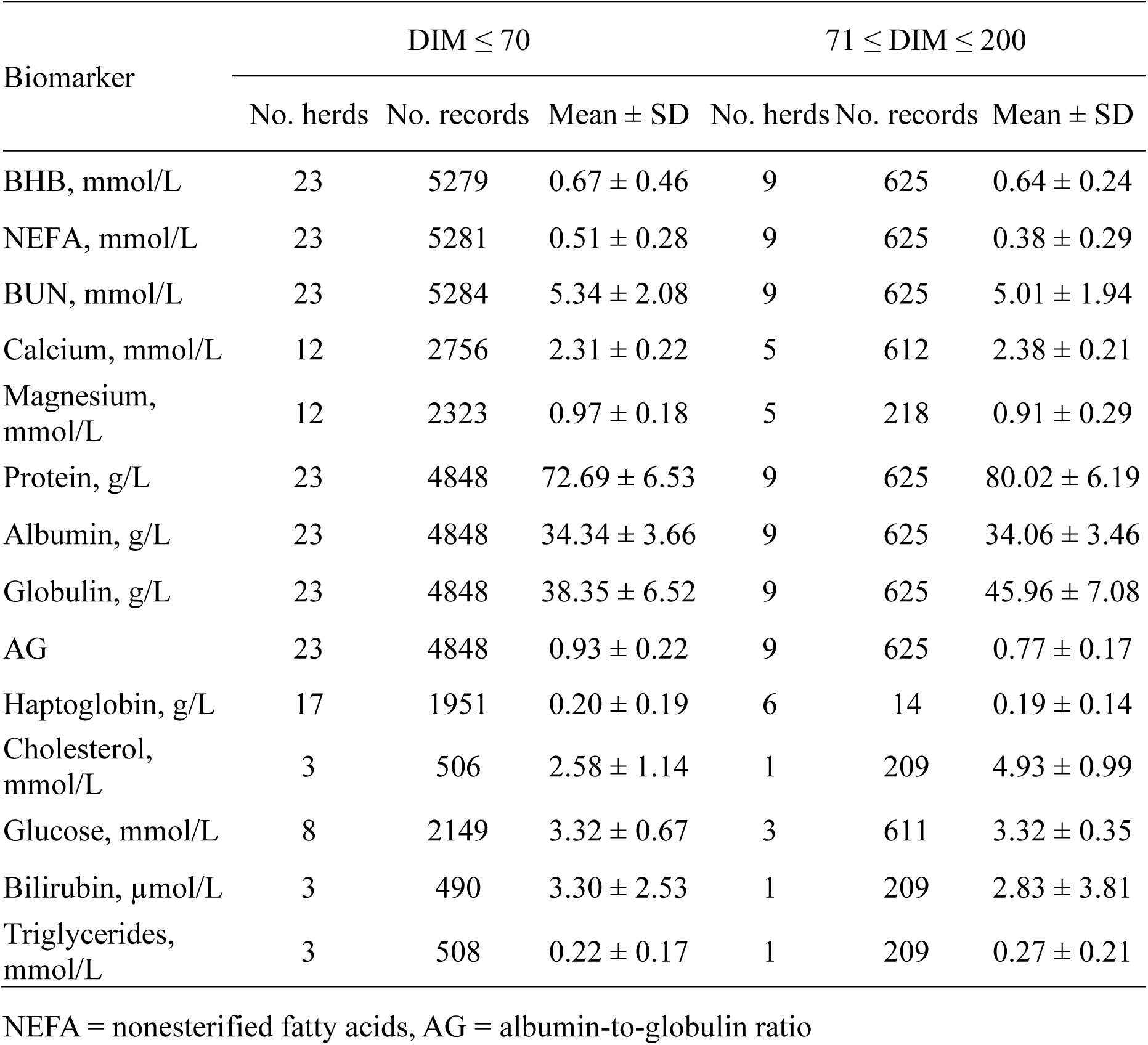
Summary statistics of serum biomarkers (mean ± SD)

Because this study focused on monitoring cow health during early lactation and given relatively few records were available from later lactation (71 ≤ DIM ≤ 200), only records collected within 70 DIM postpartum, representing approximately 90% of the data, were used for model development. Models were developed using partial least squares regression (PLSR) implemented with the caret R package (Kuhn, 2008). Model performance was assessed by nested 10-fold random cross-validation and external validation, including leave-one-herd-out validation and validation using records collected after 70 DIM.

For nested 10-fold random cross-validation, model evaluation and parameter tuning were separated using outer and inner 10-fold random cross-validation loops. In the outer loop, data were randomly partitioned into 10 folds, with each fold used once as the test set and the remaining 9 folds used for training. Within each outer training set, an inner 10-fold cross-validation was used to select the optimal number of latent variables based on root mean squared error. The model was then refitted to the full outer training set using the selected number of latent variables and evaluated on the held-out outer test fold. Predictors were centred to a mean of 0 and scaled to a variance of 1 before model fitting. This procedure was repeated 50 times, and final performance estimates were calculated as the mean of the 50 replicate means across the 10 outer folds.

For leave-one-herd-out validation, one herd at a time was excluded from model training and used as an independent validation set. Models were trained using data from the remaining herds and evaluated on the excluded herd. This process was repeated for all 23 herds. Because herd was fitted as a categorical predictor, the excluded herd was not present in the training data and therefore no herd-specific adjustment was applied to validation records from that herd. In addition, models developed using all records collected within 70 DIM were also validated using records collected after 70 DIM. This analysis was conducted to assess whether models trained using early lactation data were transferable to later lactation stages, given that previous work, such as Premi et al. (2021), suggested that several serum biomarkers remain informative beyond early lactation. Notably, only the best model identified from the nested 10-fold random cross-validation procedure was tested for external validation.

Prediction accuracy was evaluated using coefficient of determination (R^2^), root mean square error (RMSE), and ratio of performance to interquartile distance (RPIQ). According to Chang et al. (2001), a model with RPIQ > 1.89 indicates satisfactory prediction whereas RPIQ > 2.0 is considered excellent. For BHB and NEFA, predicted values were back-transformed before calculating these metrics. All statistical analyses were performed using R version 4.4.3 (R Core Team, 2026).

Summary statistics for serum biomarkers by stage of lactation are presented in Table 1. The total number of records varied among biomarkers, with BHB, NEFA, and BUN available for nearly all samples (n = 5,904 to 5,909), followed by total protein, albumin, globulin, and albumin-to-globulin ratio (n = 5,473), and glucose, calcium, and magnesium (n = 2,149 – 2,756). Cholesterol, bilirubin, and triglycerides had fewer observations (n = 699 to 717) from a small number of herds, which should be considered when interpreting the modeling results. As outlined previously, only records collected within 70 DIM were used for model development.

Mean BHB and NEFA concentrations within 70 DIM were 0.67 ± 0.46 and 0.51 ± 0.28 mmol/L, respectively. NEFA declined after 70 DIM (0.38 ± 0.29 mmol/L), whereas BHB changed only modestly (0.64 ± 0.24 mmol/L). Similar patterns have been reported, with NEFA concentrations decreasing as cows recover energy balance, whereas BHB often shows less pronounced change across lactation (Cattaneo et al., 2023). BUN concentrations were similar between early and later lactation (5.34 ± 2.08 vs. 5.01 ± 1.94 mmol/L). Although the means of NEFA and BUN within 70 DIM were comparable to previous reports (Luke et al., 2019b, Ho et al., 2021), mean BHB was higher in the present study (0.67 vs. 0.54 mmol/L). Consequently, the prevalence of hyperketonemia, defined using a threshold of ≥ 1.2 mmol/L, was higher in this study than that reported by Ho et al. (2021) (8.5 vs. 2.9%). Similarly, Brunner et al. (2018) showed a relatively high mean BHB concentration (0.7 mmol/L) and a hyperketonemia prevalence of 9.6% in 208 cows from 22 Australian farms, with blood samples collected between 2 and 21 DIM. Further investigation of the data indicated that additional blood samples collected after 2020 had considerably higher BHB concentrations than those collected between 2017 and 2020 in previous studies (0.87 vs. 0.54 mmol/L). This increased BHB appeared to be driven by high prevalence of clinical ketosis in a small number of herds.

Total protein increased from 72.69 ± 6.53 in early lactation to 80.02 ± 6.19 g/L after 70 DIM, driven primarily by globulin (38.35 ± 6.52 to 45.96 ± 7.08 g/L), whereas albumin remained stable (34.34 ± 3.66 vs. 34.06 ± 3.46 g/L). This pattern agrees with earlier observations that globulin concentrations increase postpartum as immune function recovers, whereas albumin is less influenced by lactation stage (Rowlands et al., 2009).

Macromineral concentrations showed modest stage-of-lactation differences. Calcium increased from 2.31 ± 0.22 mmol/L in early lactation to 2.38 ± 0.21 mmol/L after 70 DIM, consistent with recovery of calcium homeostasis following the periparturient period. Magnesium averaged 0.97 ± 0.18 mmol/L in early lactation and 0.91 ± 0.29 mmol/L after 70 DIM. The relatively narrow distributions of Ca and Mg reflect tight homeostatic regulation and may partly explain their limited predictability from MIR spectra.

Among indicators of hepatic function and lipid and carbohydrate metabolism, cholesterol approximately doubled from early to later lactation (2.58 ± 1.14 vs. 4.93 ± 0.99 mmol/L), whereas glucose remained unchanged (3.32 mmol/L in both periods). This strong stage-of-lactation trajectory is consistent with previous studies linking cholesterol to hepatic metabolic status and energy balance (Gross et al., 2015). Bilirubin was slightly lower after 70 DIM, and triglycerides remained low across both stages. Mean haptoglobin concentration in early lactation was 0.20 ± 0.19 g/L, but varied considerably.

Across biomarkers, model performance varied substantially between biomarkers and predictors used (Table 2). It is important to emphasize that MIR predictions are generally indirect, relying on correlations between blood biomarkers and milk components that influence the spectral signal (Eskildsen et al., 2014). In addition, MIR spectroscopy has a limited sensitivity for compounds present at very low concentrations (Dardenne et al., 2015).

**Table 2.**
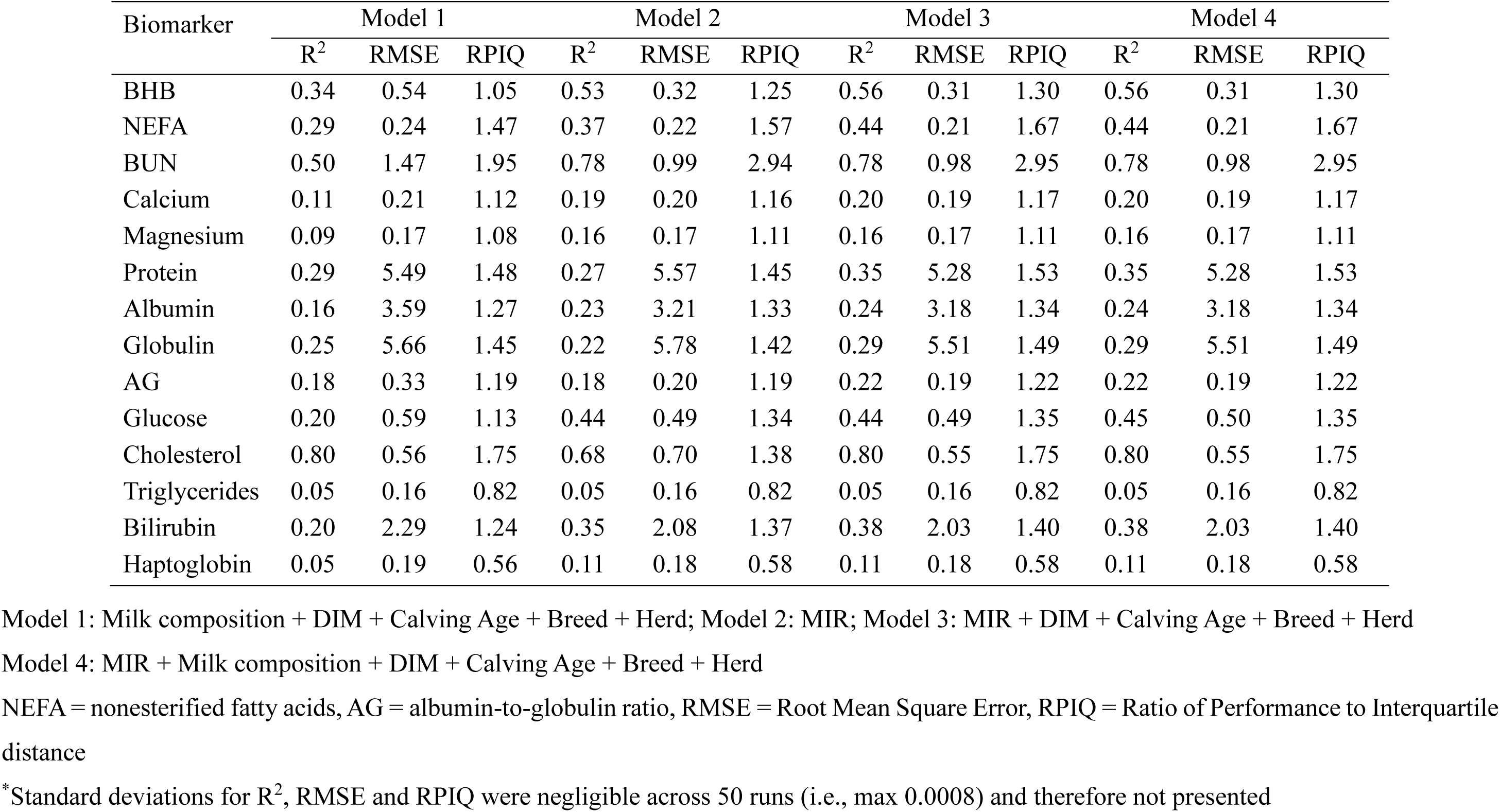
Prediction accuracy of serum biomarkers collected within 70 DIM using MIR spectra and/or on-farm variables obtained from nested 10-fold random cross validation*.

| Biomarker | Model 1 |  |  | Model 2 |  |  | Model 3 |  |  | Model 4 |  |  |
| --- | --- | --- | --- | --- | --- | --- | --- | --- | --- | --- | --- | --- |
|  | R<sup>2</sup> | RMSE | RPIQ | R<sup>2</sup> | RMSE | RPIQ | R<sup>2</sup> | RMSE | RPIQ | R<sup>2</sup> | RMSE | RPIQ |
| BHB | 0.34 | 0.54 | 1.05 | 0.53 | 0.32 | 1.25 | 0.56 | 0.31 | 1.30 | 0.56 | 0.31 | 1.30 |
| NEFA | 0.29 | 0.24 | 1.47 | 0.37 | 0.22 | 1.57 | 0.44 | 0.21 | 1.67 | 0.44 | 0.21 | 1.67 |
| BUN | 0.50 | 1.47 | 1.95 | 0.78 | 0.99 | 2.94 | 0.78 | 0.98 | 2.95 | 0.78 | 0.98 | 2.95 |
| Calcium | 0.11 | 0.21 | 1.12 | 0.19 | 0.20 | 1.16 | 0.20 | 0.19 | 1.17 | 0.20 | 0.19 | 1.17 |
| Magnesium | 0.09 | 0.17 | 1.08 | 0.16 | 0.17 | 1.11 | 0.16 | 0.17 | 1.11 | 0.16 | 0.17 | 1.11 |
| Protein | 0.29 | 5.49 | 1.48 | 0.27 | 5.57 | 1.45 | 0.35 | 5.28 | 1.53 | 0.35 | 5.28 | 1.53 |
| Albumin | 0.16 | 3.59 | 1.27 | 0.23 | 3.21 | 1.33 | 0.24 | 3.18 | 1.34 | 0.24 | 3.18 | 1.34 |
| Globulin | 0.25 | 5.66 | 1.45 | 0.22 | 5.78 | 1.42 | 0.29 | 5.51 | 1.49 | 0.29 | 5.51 | 1.49 |
| AG | 0.18 | 0.33 | 1.19 | 0.18 | 0.20 | 1.19 | 0.22 | 0.19 | 1.22 | 0.22 | 0.19 | 1.22 |
| Glucose | 0.20 | 0.59 | 1.13 | 0.44 | 0.49 | 1.34 | 0.44 | 0.49 | 1.35 | 0.45 | 0.50 | 1.35 |
| Cholesterol | 0.80 | 0.56 | 1.75 | 0.68 | 0.70 | 1.38 | 0.80 | 0.55 | 1.75 | 0.80 | 0.55 | 1.75 |
| Triglycerides | 0.05 | 0.16 | 0.82 | 0.05 | 0.16 | 0.82 | 0.05 | 0.16 | 0.82 | 0.05 | 0.16 | 0.82 |
| Bilirubin | 0.20 | 2.29 | 1.24 | 0.35 | 2.08 | 1.37 | 0.38 | 2.03 | 1.40 | 0.38 | 2.03 | 1.40 |
| Haptoglobin | 0.05 | 0.19 | 0.56 | 0.11 | 0.18 | 0.58 | 0.11 | 0.18 | 0.58 | 0.11 | 0.18 | 0.58 |

Model 1: Milk composition + DIM + Calving Age + Breed + Herd; Model 2: MIR; Model 3: MIR + DIM + Calving Age + Breed + Herd
Model 4: MIR + Milk composition + DIM + Calving Age + Breed + Herd
NEFA = nonesterified fatty acids, AG = albumin-to-globulin ratio, RMSE = Root Mean Square Error, RPIQ = Ratio of Performance to Interquartile distance
\*Standard deviations for R<sup>2</sup>, RMSE and RPIQ were negligible across 50 runs (i.e., max 0.0008) and therefore not presented

**Table 3.** Prediction accuracy (mean ± SD) of serum biomarkers collected within 70 DIM using MIR spectra, DIM, calving age, breed and herd (Model 3) obtained from external validation*.

|  | Leave-one-herd-out |  |  | Biomarker > 70 DIM |  |  |
| --- | --- | --- | --- | --- | --- | --- |
|  | R<sup>2</sup> | RMSE | RPIQ | R<sup>2</sup> | RMSE | RPIQ |
| BHB | 0.45 ± 0.17 | 0.26 ± 0.17 | 1.13 ± 0.34 | 0.004 | 0.33 | 0.95 |
| NEFA | 0.34 ± 0.17 | 0.28 ± 0.10 | 1.10 ± 0.36 | 0.003 | 0.34 | 1.29 |
| BUN | 0.58 ± 0.22 | 1.02 ± 0.49 | 2.01 ± 0.65 | 0.65 | 1.36 | 2.20 |
| Calcium | 0.19 ± 0.15 | 0.18 ± 0.05 | 1.00 ± 0.22 | 0.008 | 0.27 | 0.80 |
| Magnesium | 0.03 ± 0.04 | 0.15 ± 0.04 | 0.98 ± 0.16 | 0.20 | 0.25 | 0.99 |
| Protein | 0.19 ± 0.09 | 6.02 ± 0.97 | 1.15 ± 0.22 | 0.04 | 7.91 | 0.97 |
| Albumin | 0.11 ± 0.07 | 3.67 ± 1.37 | 1.04 ± 0.29 | 0.05 | 3.55 | 1.32 |
| Globulin | 0.16 ± 0.08 | 5.77 ± 1.05 | 1.19 ± 0.18 | 0.09 | 8.53 | 1.07 |
| AG | 0.11 ± 0.07 | 0.20 ± 0.07 | 1.13 ± 0.25 | 0.07 | 0.20 | 0.99 |
| Glucose | 0.15 ± 0.11 | 0.58 ± 0.12 | 1.01 ± 0.39 | 0.04 | 0.75 | 0.54 |
| Cholesterol | 0.13 ± 0.02 | 0.81 ± 0.37 | 0.93 ± 0.19 | 0.09 | 3.01 | 0.46 |
| Triglycerides | 0.0008 ± 0.0001 | 0.17 ± 0.03 | 0.71 ± 0.16 | 0.07 | 0.20 | 0.61 |
| Bilirubin | 0.02 ± 0.10 | 3.11 ± 0.55 | 1.11 ± 0.41 | 0.24 | 3.61 | 0.42 |
| Haptoglobin | 0.10 ± 0.07 | 0.18 ± 0.05 | 0.56 ± 0.09 | 0.08 | 0.13 | 0.56 |

NEFA = nonesterified fatty acids, AG = albumin-to-globulin ratio, RMSE = Root Mean Square Error, RPIQ = Ratio of Performance to Interquartile distance
\*Standard deviation of prediction accuracy is not available for validation using records with DIM >70

Generally, MIR spectra (Model 2) outperformed the model based only on milk composition and on-farm variables, which comprised DIM, calving age, breed, and herd (Model 1). However, the magnitude of improvement differed among biomarkers. Coefficients of determination (R^2^) increased for 9 of 14 biomarkers by 0.06 to 0.28 units, remained unchanged for AG and triglycerides, and decreased slightly for protein, globulin, and cholesterol. Combining MIR spectra with on-farm variables (Model 3) produced the highest overall performance, increasing R^2^ over MIR alone for 9 of 14 biomarkers by 0.01 to 0.12 units (4 - 32% relative improvement), with no decrease in accuracy for any biomarker. Similar findings have been reported regarding the primary contribution of MIR spectra as well as additional values of production and management variables to prediction of methane emissions and fertility of dairy cows (Shetty et al., 2017, Ho et al., 2019, Shadpour et al., 2022). In contrast, adding conventional milk composition traits (fat, protein, and lactose) to a model already including MIR spectra and on-farm variables (Model 4) did not improve accuracy. This outcome is expected because milk composition is predicted from MIR spectra. A similar finding was reported by Ho et al. (2019), who showed that incorporating MIR spectra improved predictive performance for dairy cow fertility, whereas removing milk composition traits did not reduce accuracy. Overall, these results suggest that combining MIR spectra with on-farm variables produced the most effective model for predicting serum biomarkers in dairy cows. Therefore, external validation was conducted using Model 3 only.

Regarding individual biomarkers, BHB and NEFA were moderately predicted, with R² values of 0.56 and 0.44, RMSE of 0.31 and 0.21 mmol/L, and RPIQ of 1.30 and 1.67, respectively. BUN was the most accurately predicted biomarker (R² = 0.78, RPIQ = 2.95). These results are consistent with previous studies reporting that BHB, NEFA, and BUN are among the most predictable blood biomarkers from milk MIR spectra (Benedet et al., 2019, Bonfatti et al., 2019, Ho et al., 2021). Luke et al. (2019b), for example, reported R^2^ values of 0.48, 0.61, and 0.90 for serum BHB, NEFA, and BUN, respectively, while these were 0.63, 0.52, and 0.58 in Benedet et al. (2019). It is important to reiterate that the data used in Luke et al. (2019b) and Ho et al. (2021) were part of that analyzed in this study.

Protein fractions (albumin, globulin and AG), macrominerals (Ca and Mg), and haptoglobin showed only low to moderate prediction accuracy (R^2^ = 0.11 – 0.35). Glucose and bilirubin were moderately predicted (R² = 0.44 and 0.38, respectively), whereas triglycerides showed negligible predictability (R² = 0.05). The low predictability of triglycerides is consistent with the results of Benedet et al. (2019). Cholesterol achieved relatively high accuracy (R² = 0.80, RPIQ = 1.75). However, a further inspection of the scatter plot between predicted and observed cholesterol revealed two distinct clusters corresponding to low and high cholesterol concentrations before and after 30 DIM, respectively, which appeared to largely drive the high overall R^2^. Within each cluster, the R² was approximately 0.27, indicating considerably weaker within-stage predictive ability. This suggests that the model might have captured broad lactation-stage differences rather than accurately predicting individual cow variation in cholesterol concentration, despite DIM already being included as a predictor.

Compared with the literature, our glucose, cholesterol and triglycerides prediction accuracy was similar to that reported by Benedet et al. (2019), with R^2^ = 0.20, 0.16, and 0.39 respectively. In contrast, the prediction accuracy for protein fractions and macrominerals was lower than that reported in some other studies. For example, Soyeurt et al. (2009) reported cross-validation R² values of 0.80, 0.50, 0.70, 0.79, and 0.23 for milk Ca, Mg, Na, P, and K, respectively. However, those results were based on relatively small calibration sets, with only 31 to 57 samples used for mineral reference analyses, which may limit direct comparison with the present study. Giannuzzi et al. (2023) investigated MIR-based prediction of a broader panel of 29 blood biomarkers using machine-learning methods. They reported comparatively good prediction accuracy (R^2^ = 0.34 - 0.78) for several biomarkers related to energy metabolism, liver function and hepatic damage, inflammation and innate immunity, and some protein fractions, whereas prediction accuracy for oxidative stress indicators and minerals was generally more variable (R^2^ = 0.20 - 0.78).

Although random cross-validation is commonly used to assess the performance of MIR prediction equations, it often provides over-optimistic results, especially when records from the same herds or same test date are represented in both training and validation sets. In our dataset, there were 5,936 blood samples from 4,442 cows, which means that some cows contributed more than one record. Wang and Bovenhuis (2019) reported a substantial decline in the prediction accuracy (R^2^) of methane emissions using MIR from 0.49 to 0.01 on random cross-validation and leave-one-herd-out validation, respectively. This reduction reflects the difficulty of transferring MIR equations across herds differing in management, feeding system or breed composition. This emphasizes the importance of externally validating the model before practical implementation. In the current study, model transferability was evaluated using both leave-one-herd-out validation and validation in cows sampled after 70 DIM (treated as a single data block), using models trained exclusively on records collected within 70 DIM.

External validation showed clear differences among biomarkers in model robustness across herds and lactation stages. Blood urea nitrogen had the strongest and most consistent external performance, with leave-one-herd-out R² = 0.58 ± 0.22, RMSE = 1.02 ± 0.49 mmol/L, and RPIQ = 2.01 ± 0.65. Performance remained comparable in cows sampled after 70 DIM (R² = 0.65; RPIQ = 2.20). However, the relatively large standard deviation in leave-one-herd-out R^2^ indicates that more data, especially across different feeding and management systems should be collected to improve the robustness of the model (Pralle and White, 2020). BHB and NEFA showed moderate leave-one-herd-out performance, with average R² values of 0.45 ± 0.17 and 0.34 ± 0.17, respectively. However, their performance was strongly stage-dependent, declining to near zero when applied to cows sampled after 70 DIM (R^2^ = 0.004 and 0.003, respectively). This suggests that these equations captured the early-lactation metabolic signal associated with negative energy balance, lipid mobilisation, and ketogenesis, but were not transferable beyond this biological window. Using leave-one-herd-out validation, Gruber et al. (2026) reported MIR-BHB prediction accuracies of R^2^ = 0.18 with PLSR and R^2^ = 0.25 with support vector machine or artificial neural network models.

Hepatic, lipid, carbohydrate-related, and inflammatory biomarkers showed poor transferability overall. Glucose had low accuracy in both leave-one-herd-out and >70 DIM validation (R^2^ = 0.15 and 0.04, respectively), while bilirubin, triglycerides, and haptoglobin were not reliably predicted. Haptoglobin showed poor external validity (leave-one-herd-out R^2^ = 0.10 and >70 DIM R^2^ = 0.08). Cholesterol provided the clearest example of overestimation by random cross-validation: despite high random cross-validation accuracy (R^2^ = 0.80), external validation performance was poor (leave-one-herd-out R^2^ = 0.13 and >70 DIM R^2^ = 0.09), suggesting that the apparent accuracy could be driven by DIM-related structure rather than a robust MIR-serum cholesterol relationship. However, it should be noted that the number of herds available for validation differed among traits: bilirubin, cholesterol, and triglycerides were assessed in only three herds; glucose in eight herds; and the remaining biomarkers in 12 to 23 herds. Additional data from more herds would be required for a robust validation.

Overall, external validation confirmed that the MIR-based model for BUN had the most robust and transferable predictive performance, whereas models for BHB and NEFA showed moderate leave-one-herd-out performance but were limited to early lactation. Potential applications of these predictions could include monitoring and genetic evaluation for nitrogen efficiency, ketosis, or energy deficit. In contrast, models for the remaining biomarkers had low external prediction accuracy.

## Notes

This study was undertaken as part of the DairyBio program, which is jointly funded by Dairy Australia (Melbourne, Australia), the Gardiner Foundation (Melbourne, Australia), and Agriculture Victoria (Melbourne, Australia). The authors thank all farmers who participated in this project, as well as the staff at Ellinbank SmartFarm (Victoria, Australia) for their assistance with animal feeding and husbandry. The authors also thank Thuy Nguyen, Tim Sarget, and Gert Nieuwhof of DataGene (Bundoora, Australia) for providing MIR and cow data. In addition, the authors gratefully acknowledge the field team - Anastasiia Kudriashova, Boris Sepulveda, Brett Mason, Christian Krill, Christy Vander Jagt, Doris Ram, Ee Cheng Ooi, Fazel Almasi, Irene van den Berg, Jessica McArt, Jianghui Wang, Jo Newton, Joanne Bui, Laura Jensen, Priyanka Reddy, Ren Hatherley, and Wenjiao Wang - for assisting with blood sampling. The authors also thank Mitch Crawford of Rochester Veterinary Practice, Bernie Baxter of Dairy Australia, and Laura Calder of DataGene for their assistance with farmer recruitment.

### Ethics statement

All animal procedures were conducted in accordance with the Australian Code for the Care and Use of Animals for Scientific Purposes (NHMRC, 2013). Approval was obtained from the DEECA Agricultural Research and Extension Animal Ethics Committee for samples collected after April 2020 and from the Tasmanian Department of Primary Industries, Parks, Water and Environment for the 2018 samples collected from 2 Tasmanian herds.

